# Mechanosensitive calcium signaling in response to cell shape changes promotes epithelial tight junction remodeling by activating RhoA

**DOI:** 10.1101/2021.05.18.444663

**Authors:** Saranyaraajan Varadarajan, Rachel E. Stephenson, Eileen R. Misterovich, Jessica L. Wu, Ivan S. Erofeev, Andrew B. Goryachev, Ann L. Miller

## Abstract

Epithelia maintain an effective barrier by actively remodeling cell-cell junctions in response to mechanical stimuli. Cells often respond to mechanical stress through activation of RhoA and dynamic remodeling of actomyosin. Previously, we found that local leaks in the epithelial tight junction barrier are rapidly repaired by localized, transient activation of RhoA, a process we termed “Rho flares”, but how Rho flares are initiated remains unknown. Here, we discovered that intracellular calcium flashes occur in *Xenopus laevis* embryonic epithelial cells undergoing rapid remodeling of tight junctions via activation of Rho flares. Calcium flashes originate at the site of leaks and propagate into the cell. Depletion of intracellular calcium or inhibition of mechanosensitive calcium channels (MSC) reduced the amplitude of calcium flashes and diminished the activation of Rho flares. Furthermore, MSC-dependent calcium influx was necessary to maintain global barrier function by regulating repair of local tight junction proteins through efficient contraction of junctions. Collectively, we propose that MSC-dependent calcium flashes are an important mechanism allowing epithelial cells to sense and respond to local leaks induced by mechanical stimuli.

## Introduction

The ability of epithelial tissues to establish and maintain a selective paracellular barrier is crucial for development of multicellular organisms and proper function of vital organs (Ivanov et al., 2010; Luissint et al., 2016; Marchiando et al., 2010; Moriwaki et al., 2007). The dynamic nature of epithelial tissues during development and organ homeostasis requires cells to change shape and remodel their cell-cell junctions during tissue-generated forces including cell division, extrusion, and intercalation (Gudipaty and Rosenblatt, 2016; Guillot and Lecuit, 2013; Varadarajan et al., 2019). Remarkably, epithelia are able to maintain barrier function despite this remodeling (Higashi et al., 2016; Rosenblatt et al., 2001; Stephenson et al., 2019). However, the mechanisms by which epithelial cells remodel their cell-cell junctions without compromising barrier function is not fully understood.

In vertebrates, epithelial barrier function is regulated by the apical junctional complex, including tight junctions (TJs) and adherens junctions (AJs) (Zihni et al., 2016). Claudins and Occludin, transmembrane TJ proteins, interact with their counterparts on neighboring cells and polymerize within the membrane to form a network of TJ strands that selectively regulate the passage of small ions and macromolecules between the cells (Claude and Goodenough, 1973; Furuse et al., 1998; Staehelin et al., 1969). TJ strands are connected to the underlying actomyosin array by the zonula occludens (ZO) family of scaffolding proteins, thereby coupling barrier function to actomyosin-mediated cells shape changes (Fanning et al., 1998; Furuse et al., 1998; Itoh et al., 1999; Van Itallie et al., 2017). Although it is known that mechanical stress leads to a global increase in permeability to macromolecules (Cavanaugh et al., 2006; Cohen et al., 2008; Samak et al., 2010), much less is known about how cells respond to these leaks.

We recently reported the occurrence of short-lived leaks associated with cell-generated mechanical forces in the developing *Xenopus* epithelium (Stephenson et al., 2019). During *Xenopus* gastrulation, the epithelial monolayer covering the animal cap of the embryo undergoes frequent cell divisions and morphogenetic movements, requiring elongation of cell-cell junctions to accommodate cell shape changes. Leaks were associated with elongating junctions and happened at sites where TJ proteins were locally reduced. Leaks were dynamically repaired by localized, short-lived activations of the small GTPase RhoA, which we termed “Rho flares” (Stephenson et al., 2019). Rho flares are associated with local membrane protrusion at the site of leaks, suggesting that RhoA may be activated by a membrane tension-mediated mechanosensitive pathway.

Calcium signaling plays a key role in transducing mechanical forces into biochemical signals during cell shape changes (Christodoulou and Skourides, 2015; Sahu et al., 2017). Calcium signaling varies from long-range calcium waves to subcellular calcium transients, both of which are controlled precisely in space and time to modulate cytoskeleton-mediated cell shape changes. Calcium signaling is required for local RhoA activation during mechanical events including wound healing (Benink and Bement, 2005; Clark et al., 2009), cell migration (Pardo-Pastor et al., 2018), and dendritic spine enlargement (Murakoshi et al., 2011), suggesting the possibility of calcium-mediated RhoA activation during TJ remodeling.

Elevation of intracellular calcium following a mechanical stimulus is mediated by influx of extracellular calcium through mechanosensitive calcium channels (MSCs) in the plasma membrane or calcium release from the endoplasmic reticulum (ER) (Shao et al., 2015). Plasma membrane localized MSCs – including the Piezo and Transient Receptor Potential (TRP) families – allow calcium influx in response to local and/or global changes in membrane tension or curvature (Coste et al., 2010; Gudipaty et al., 2017; Liu and Montell, 2015; Miyamoto et al., 2014; Mochizuki et al., 2009; Shi et al., 2018). For example, during cell migration, MSCs mediate transient calcium influx at lamellipodia and focal adhesions to guide the direction of migration (Ellefsen et al., 2019; Wei et al., 2009). However, the role of MSCs in force-dependent remodeling of TJs has not been elucidated.

Here, we investigate whether calcium signaling and MSCs are involved in TJ remodeling using live imaging of the gastrula-stage *Xenopus* epithelium. We show that a local intracellular calcium increase follows paracellular leaks in response to cell-cell junction elongation. We demonstrate that this intracellular calcium increase is required for the Rho flare-mediated TJ repair pathway and is dependent on MSCs. Finally, we show that MSC-mediated calcium influx regulates local junction contraction through robust Rho activation and is required to repair and maintain an intact barrier during cell shape changes.

## Results

### Epithelial barrier leaks induce a local intracellular calcium increase

Intracellular calcium signaling has been implicated in TJ biogenesis, establishment of barrier function, and actomyosin-mediated cell shape changes (Christodoulou and Skourides, 2015; Gonzalez-Mariscal et al., 1990; Nigam et al., 1992; Suzuki et al., 2017). Thus, we hypothesized that intracellular calcium signaling might be involved in Rho flare-mediated TJ repair of local leaks. To test this idea, we performed Zinc-based Ultrasensitive Microscopic Barrier Assay (ZnUMBA (Stephenson et al., 2019)) in gastrula-stage *Xenopus* embryos (Nieuwkoop and Faber stage 10.5-12) expressing probes for intracellular calcium and active Rho (tagBFP-C2 (Yu and Bement, 2007) and mCherry-2xrGBD (Benink and Bement, 2005; Davenport et al., 2016)). The C2 domain of PKCβ, when bound to calcium, is recruited from the cytoplasm to the plasma membrane, and is capable of detecting both local and global calcium increases. High-speed live imaging demonstrated that calcium increases locally at sites of leaks (Fig. 1 A). FluoZin-3 intensity increased prior to the local calcium increase at the site of Rho flares, indicating a leak, (Fig. 1 B) and returned to baseline following the increase in calcium and active Rho, indicating restoration of barrier function (Fig. 1 B and Video 1).

**Figure 1:**
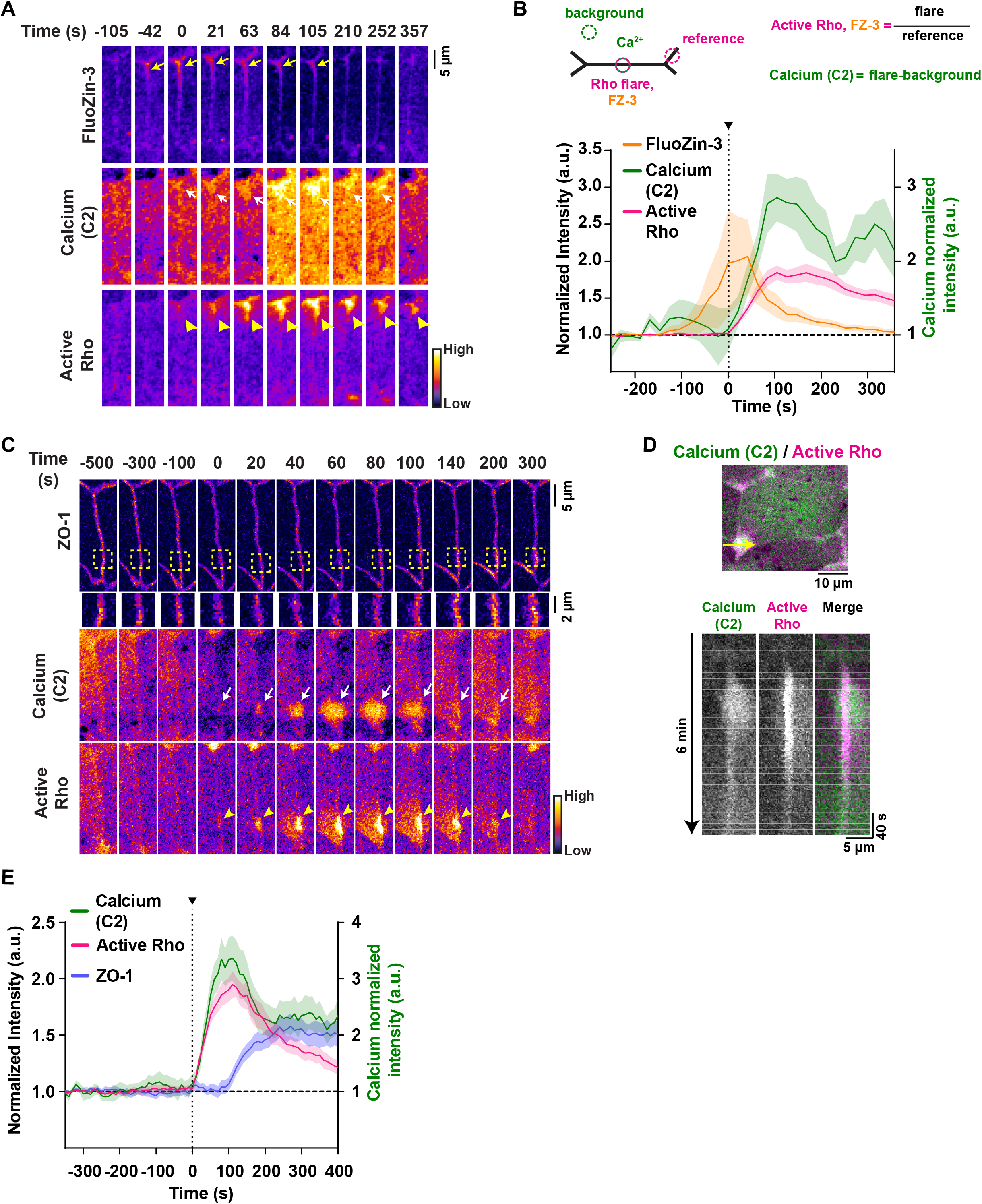
Epithelial paracellular leaks induce a local intracellular calcium increase. (A) Time lapse images (Fire LUT) of FluoZin-3 dye, membrane calcium probe (tagBFP-PKC β-C2), and active Rho (mCherry-2xrGBD). Calcium increase (white arrows) follows a leak indicated by increase in FluoZin-3 fluorescence (yellow arrows) at the site of Rho flare (yellow arrowheads). Time 0 s represents start of Rho flare. (B) Quantification of experiments shown in A. Top: Schematic showing regions of interest (ROIs) and used to quantify Rho flares, calcium (C2), and FluoZin-3. Bottom: Graph of mean normalized intensity shows the leak (FluoZin-3) precedes the increases in local calcium and active Rho. Shaded region represents S.E.M. n= 18 flares, 10 embryos, 6 experiments. (C) Time lapse images (FIRE LUT) of BFP-ZO-1, membrane calcium probe (mNeon-PKC β-C2), and active Rho (mCherry-2xrGBD). Local calcium increase (white arrows) is spatially localized to the site of ZO-1 decrease (yellow boxed region) and Rho flare (yellow arrowheads). (D) Top: Cell view of an embryo expressing membrane calcium probe (mNeon-PKCβ-C2, green) and active Rho probe (mCherry-2xrGBD, magenta). The yellow arrow indicates the 5-pixel wide region used to generate the kymograph. Bottom: Kymograph shows that calcium increase originates at the junction and spreads. (E) Quantification of experiments shown in C. Graph of mean normalized intensity shows that local calcium increases simultaneously with the increase in active Rho. Shaded region represents S.E.M. n= 19 flares, 15 embryos, 8 experiments.

To better evaluate the spatial origin of the calcium increase at Rho flares, we generated a kymograph at the site of the Rho flare. The kymograph shows that the intracellular calcium increase originates at the plasma membrane and spreads asymmetrically into the cells adjacent to the membrane (Fig. 1 D). Asymmetric intracellular calcium increase is proportional to the active Rho intensity in each of the cells adjacent to the membrane (Fig. 1, C-D and Video 2). Because Rho flares precede TJ reinforcement (Stephenson et al., 2019), we next examined whether the local calcium increase precedes ZO-1 reinforcement. Indeed, quantification revealed that the local calcium increase precedes ZO-1 reinforcement (Fig. 1, C and E). Together, these results show that the local calcium increase originates at sites of leaks and is followed by ZO-1 reinforcement.

### Intracellular calcium flash precedes Rho flares during local ZO-1 reinforcement

To better characterize the temporal dynamics of the intracellular calcium increase, we imaged GCaMP6m (Chen et al., 2013), a genetically encoded calcium probe with faster kinetics but reduced spatial sensitivity compared to C2. We detected a transient increase in calcium in cells adjacent to the junction with the Rho flare (Fig. 2 A) with a duration of 72.72 ± 9.13 s (Fig. 2 B), hereafter referred to as a “calcium flash”. Calcium flashes associated with Rho flares differ from calcium waves previously described during *Xenopus* gastrulation because they are restricted to the cells neighboring the junction exhibiting a Rho flare (Fig. S1, A-B and Video 3). In contrast, calcium waves originate in 2-4 cells and propagate to neighboring cells within minutes (Wallingford et al., 2001) (Fig. S1, C-C’ and Video 3).

**Figure 2:**
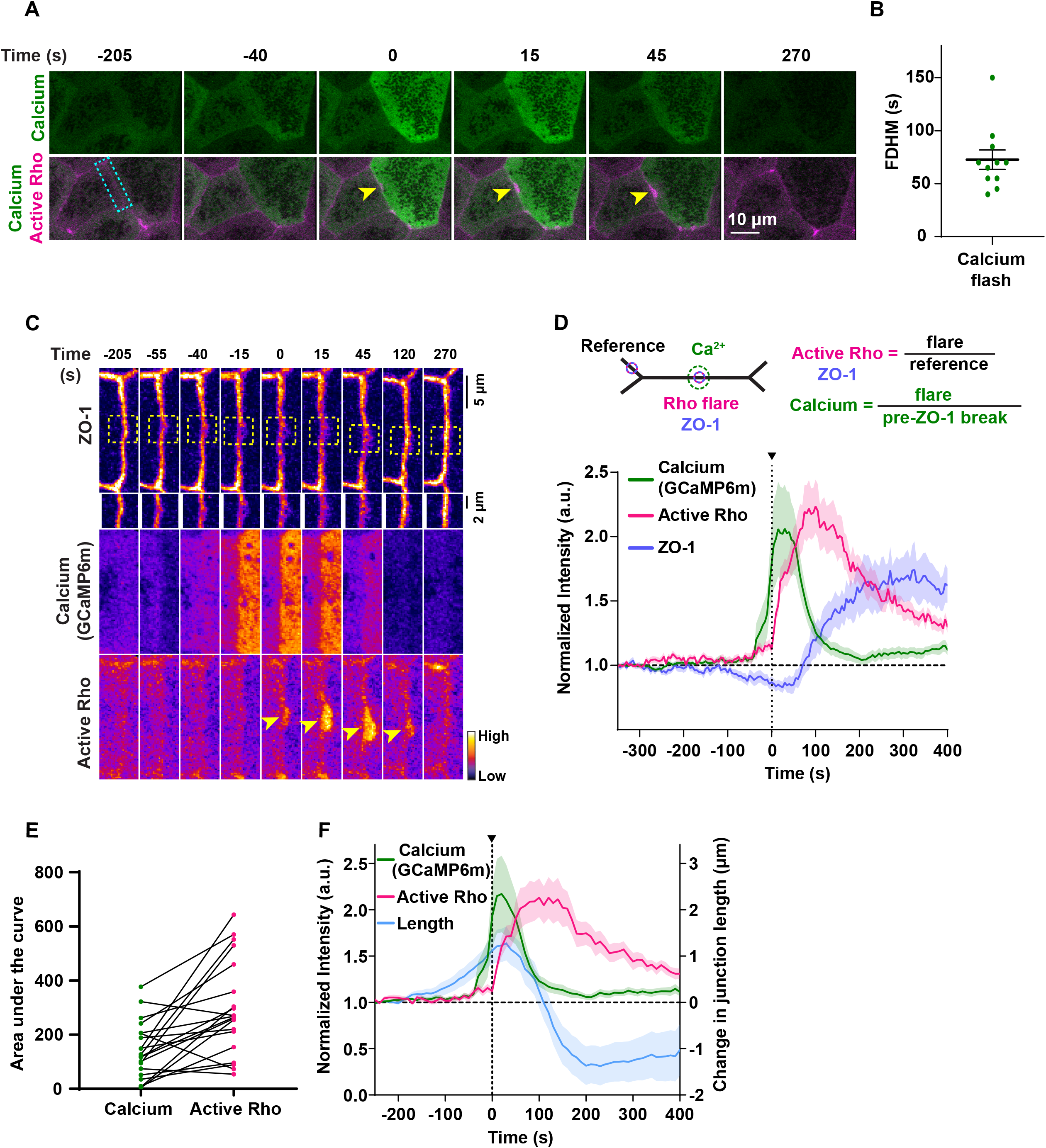
Intracellular calcium increase precedes Rho flares during local ZO-1 reinforcement. (A) Time lapse images of calcium (GCaMP6m, green) and active Rho (mCherry-2xrGBD, magenta). Calcium flash occurs in cells adjacent to the junction with the Rho flare (yellow arrowheads). Time 0 s represents start of Rho flare. (B) Dot plot of calcium flash duration (full duration at half maximum, FDHM). Error bar represents mean ± S.E.M. n= 11 flashes, 11 embryos, 10 experiments. (C) Montage of the junction indicated by the blue box in A (FIRE LUT). A calcium flash (GCaMP6m) follows a local loss of ZO-1 (BFP-ZO-1, yellow boxed region, enlarged below) and precedes the Rho flare (mCherry-2xrGBD, yellow arrowhead). (D) Quantification of experiments shown in C. Top: Schematic showing regions of interest (ROIs) used to quantify Rho flares, calcium, and ZO-1. Bottom: Graph showing calcium intensity increase follows a decrease in ZO-1 and precedes an increase in active Rho. Shaded region represents S.E.M. n= 20 flares, 15 embryos, 12 experiments. (E) Dot plot of area under the curve for calcium (green) and active Rho (magenta) curves in D showing the correlation (solid black lines) of calcium intensity and corresponding Rho flare intensity. n= 20 flares, 15 embryos, 12 experiments. (F) Graph showing calcium flash (GCaMP6m) follows junction elongation and precedes Rho-mediated junction contraction and stabilization of length. Shaded region represents S.E.M. n= 17 flares, 14 embryos, 11 experiments.

When observed with GCaMP6m, calcium flashes occur after the decrease in ZO-1 and precede Rho flares by ∼40-50 s (Fig. 2, C and D). The intensity of GCaMP6m decreases as reinstatement of ZO-1 is initiated (Fig. 2 D and Video 4). As Rho flares vary in size and amplitude, we hypothesized that the amplitude of the calcium flash scales with Rho flare amplitude. To evaluate this idea, we plotted the area under the curve for calcium and active Rho for individual flares. Indeed, we observed that higher intensity calcium flashes generally correlated with higher intensity, sustained Rho flares (Fig. 2 E and Video 5).

To evaluate junction contraction in relation to the calcium flash, we measured the vertex-to-vertex junction distance of the cell-cell junctions where Rho flares occur over time. We found that junction elongation preceded the calcium flash, and a rapid contraction of the junction immediately followed the calcium flash and Rho flare (blue trace, Fig. 2 F). Of note, we observed that the initiation of junction contraction occurs during the peak of the calcium flash, and the stabilization of junction length occurs as the calcium level returns to baseline. Taken together, our results support a role for calcium flashes in promoting Rho flare-mediated TJ reinforcement through contraction and stabilization of cell-cell junctions.

### Intracellular calcium flash is required for Rho flare activation and ZO-1 reinforcement

Calcium is required for Rho activation during local cell shape changes including epithelial wound healing, cell migration, and dendritic spine enlargement (Clark et al., 2009; Murakoshi et al., 2011; Pardo-Pastor et al., 2018). However, whether calcium is required for local Rho activation and ZO-1 reinforcement at leaks remains unknown. To test this question, we chelated intracellular calcium using BAPTA-AM, a cell-permeable calcium chelator that chelates intracellular calcium without disrupting calcium-dependent cadherin adhesion. BAPTA-AM alone treatment led to an increase in the frequency of calcium waves (Fig. S2 A), which disrupted our ability to observe calcium flashes. Calcium waves are dependent on ER-mediated calcium release (Wallingford et al., 2001), so we blocked them using the IP_3_ channel blocker, 2-APB, in addition to chelating intracellular calcium with BAPTA-AM (Fig. 3 A). Upon treatment with BAPTA-AM + 2-APB (Fig. 3 A’), the intensity of calcium flashes was reduced compared to vehicle (DMSO) (Fig. 3, B-D’ and Video 6). Active Rho was also significantly reduced at sites of ZO-1 loss when intracellular calcium was chelated (Fig. 3, B, C, E, and E’, and Video 6). Finally, we observed that ZO-1 breaks were more severe, lasted longer, and failed to be reinstated in intracellular calcium chelated embryos compared to vehicle controls (Fig. 3, B, C, F and F’, and Video 6). Together, our results suggest that intracellular calcium increase is required for both Rho flare activation and efficient ZO-1 reinforcement.

**Figure 3:**
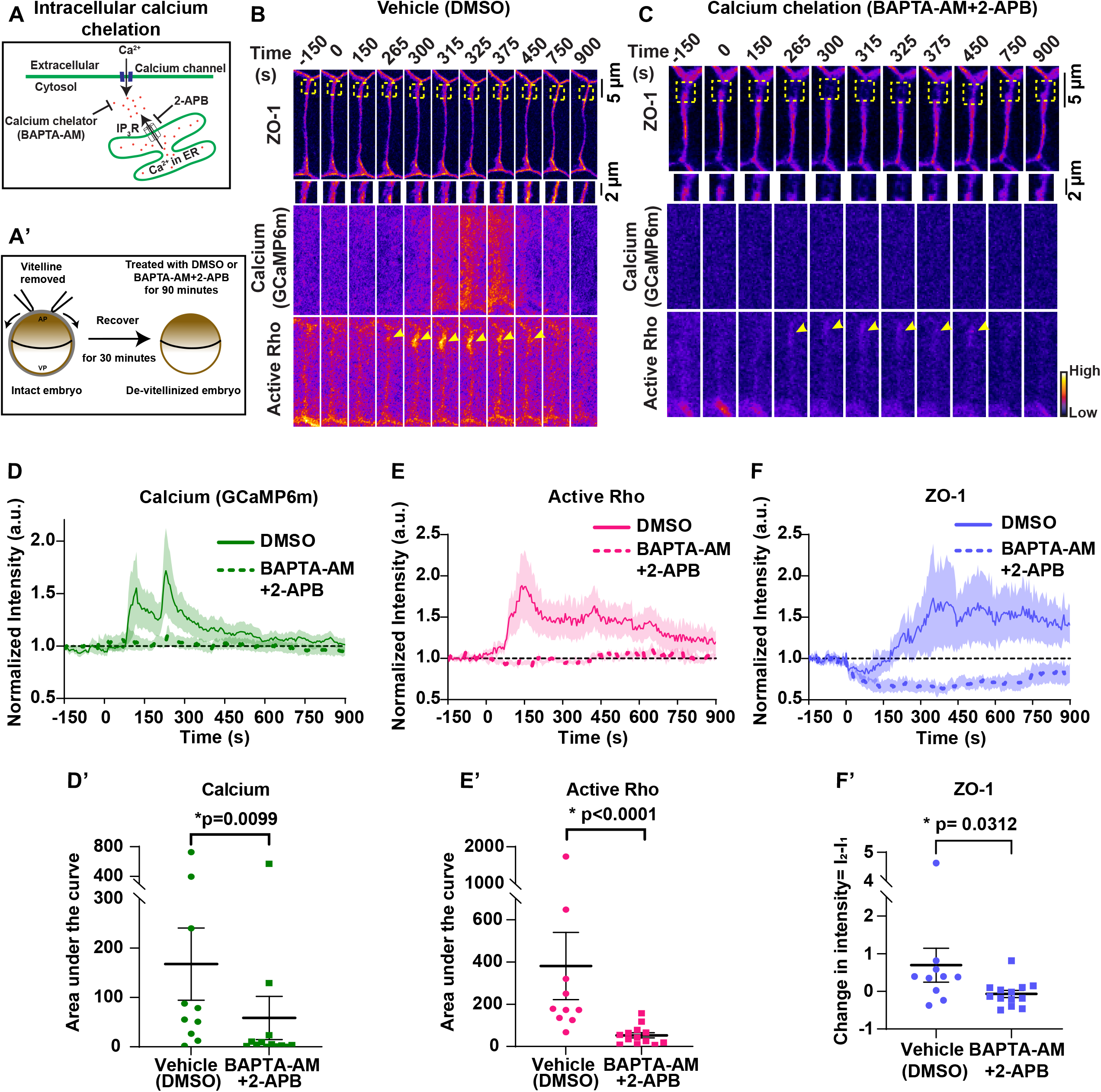
Intracellular calcium flash is required for Rho flare activation and ZO-1 reinforcement. (A) Schematic of cytosolic calcium chelation using BAPTA-AM and blocking IP_3_R-mediated calcium release from the ER using 2-APB. (A’) Schematic showing BAPTA-AM and 2-APB treatment after removal of vitelline envelope. (B-C) Time-lapse images (FIRE LUT) of BFP-ZO-1, calcium (GCaMP6m), and active Rho (mCherry-2xrGBD). Embryos were treated with 0.5% DMSO (vehicle, B) or 20 µM BAPTA-AM and 100 µM 2-APB (calcium chelation, C). Calcium chelation resulted in more severe ZO-1 breaks (yellow boxes, enlarged below) and decreased Rho activity (yellow arrowheads) at the site of ZO-1 loss. Time 0 s represents the start of ZO-1 decrease. (D-F) Graphs of mean normalized intensity of calcium (GCaMP6m, D), active Rho (E) and ZO-1 (F) at the site of ZO-1 loss over time in vehicle (DMSO, solid lines) or calcium chelation (BAPTA-AM+2-APB, dashed lines) embryos. Shaded region represents S.E.M. DMSO: n= 10 flares, 7 embryos, 7 experiments; BAPTA-AM+2-APB: n=13 flares, 8 embryos, 8 experiments. (D’-F’) Area under the curve for calcium (D’) and active Rho (E’) was calculated from D and E, respectively. Scatter plot of change in ZO-1 intensity (F’) was calculated from F. I_1_ and I_2_ represent average intensity of ZO-1 from individual traces from time 0-50 s and 400-450 s, respectively. Error bars represent mean ± S.E.M.; significance calculated using Mann-Whitney test.

### Mechanosensitive calcium channel-dependent calcium influx is required for Rho-mediated reinforcement of ZO-1

Plasma membrane localized MSCs play an important role in mediating local calcium influx in response to changes in local membrane tension (Canales et al., 2019; Ellefsen et al., 2019). Rho flares occur following junction elongation (Fig. 2 F) and loss of ZO-1 (Fig. 2 D), a scaffolding protein that connects TJ transmembrane proteins to the underlying actin cytoskeleton, suggesting a possible change in membrane tension due to detachment of membrane from the underlying actin cytoskeleton. Furthermore, Rho flares are associated with apical plasma membrane deformation (Stephenson et al., 2019) and a local calcium increase that originates at the plasma membrane (Fig. 1 D). Together, these observations suggest that calcium influx may be mediated by MSCs. First, we evaluated the temporal association of membrane protrusion (tagBFP-membrane) and Rho flares (mCherry-2xrGBD). The kymograph at the site of a Rho flare showed that membrane protrusion (blue, Fig. 4 A) expands with a similar timing to the increase in Rho activation (magenta, Fig. 4 A) supporting a mechanosensation-based mechanism. Therefore, to test if MSCs are responsible for the calcium flash preceding Rho flares, we blocked MSCs using Grammostola Mechanotoxin #4 (GsMTx4), a peptide from spider venom. GsMTx4 decreases the sensitivity of MSCs to changes in membrane tension, thereby blocking calcium influx mediated by Piezo1 and TRP channels (Fig. 4 B) (Bae et al., 2011; Bowman et al., 2007; Gnanasambandam et al., 2017). MSC-mediated calcium transients are required for tissue homeostasis and proper development (Hunter et al., 2014). Therefore, we used GsMTx4 at a low concentration such that embryos successfully develop to gastrula stage, and the overall morphology of cell-cell junctions is not altered. With GsMTx4-treatment, we observed a significant decrease in the calcium flash associated with Rho flares (Fig. 4, C-E and E’), indicating that the calcium flashes preceding Rho flares are mediated by MSCs. Further, with GsMTx4 treatment, the amplitude and duration of Rho flares were significantly reduced compared to vehicle (water) (Fig. 4, C-D, F, and F’). Although both vehicle- and GsMTx4-treated embryos showed an initial increase in active Rho at the site of ZO-1 loss, Rho flares were lower in intensity and short-lived in GsMTx4-treated embryos (Fig. 4 F). In addition, by measuring the normalized intensity of ZO-1 over time, we found that the amount of ZO-1 loss prior to Rho activation was comparable between vehicle- and GsMTx4-treated embryos (I_1_, Fig. 4 G). However, treatment with GsMTx4 significantly reduced ZO-1 reinforcement compared to vehicle controls (Fig. 4, C-D, G, and G’). Of note, GsMTx4 treatment did not alter active RhoA accumulation at the contractile ring in dividing cells (Fig. S2 B), or steady state levels of active RhoA (Fig. S2, C-C’), or the downstream targets of active RhoA (Fig. S3, D-E) at cell-cell junctions. Together, these findings demonstrate that MSC-mediated calcium influx is required for both robust Rho flare activation and efficient reinforcement of ZO-1.

**Figure 4:**
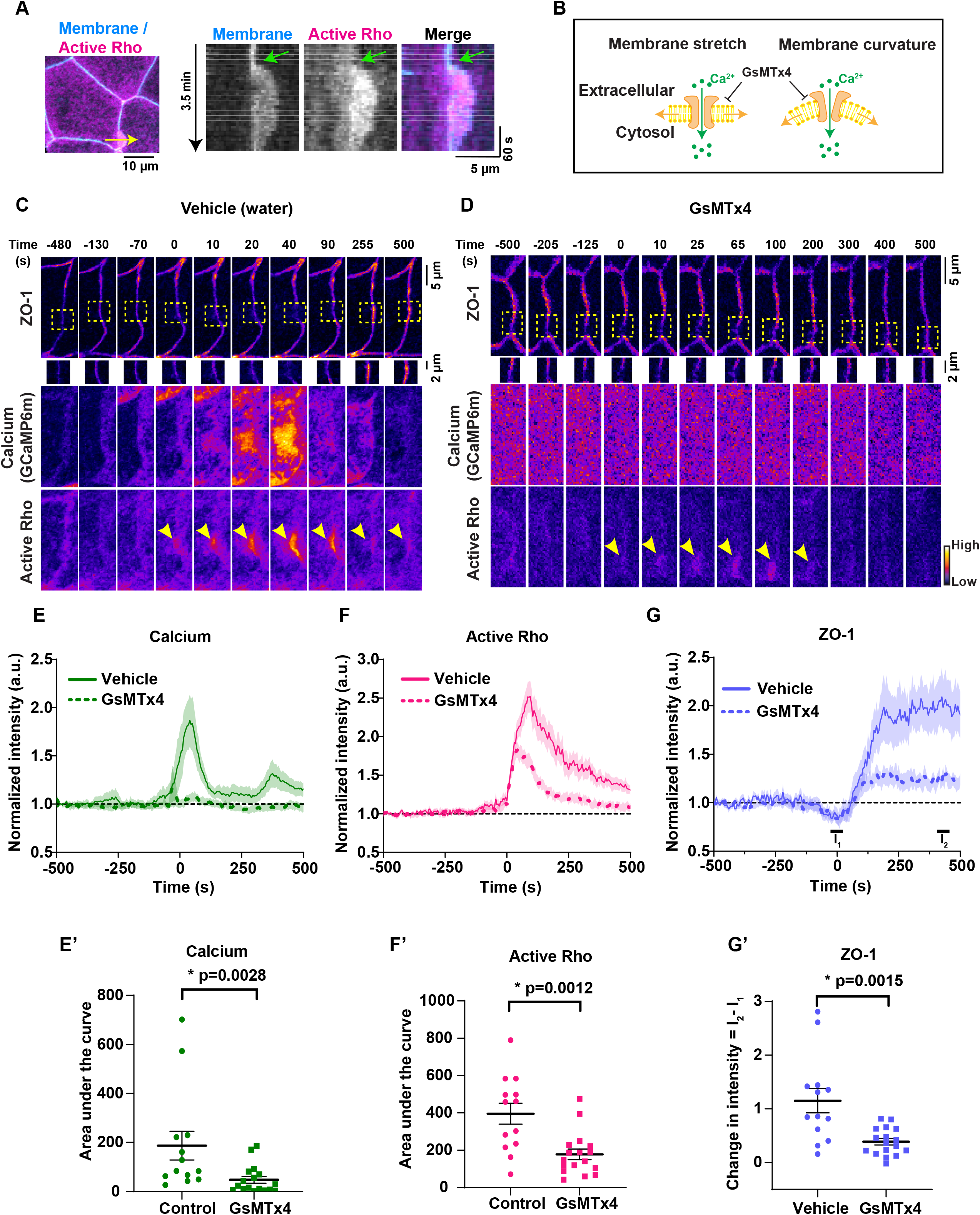
Mechanosensitive calcium channel-dependent calcium influx is required for Rho-mediated reinforcement of ZO-1. (A) Left: Cell view of an embryo expressing membrane probe (membrane-tagBFP, blue) and active Rho probe (mCherry-2xrGBD, magenta). Yellow arrow indicates the 2-pixel wide region used to generate the kymograph. Right: Kymograph shows that membrane protrusion increases with Rho flare. Green arrow indicates the start of membrane protrusion and Rho flare. (B) Schematic showing that GsMTx4 inhibits MSC-mediated calcium influx in response to changes in membrane tension induced by stretch or curvature. (C-D) Time-lapse images (FIRE LUT) of BFP-ZO-1, calcium (GCaMP6m), and active Rho (mCherry-2xrGBD). Embryos were treated with water (vehicle, C) or 12.5 µM GsMTx4 (MSC inhibitor, D). MSC inhibition resulted in a decrease in calcium influx, short duration, low intensity Rho flares (yellow arrowheads), and decreased ZO-1 reinforcement at the site of ZO-1 loss (yellow boxes, enlarged below). Time 0 s represents the start of Rho flares. (E-G) Graphs of mean normalized intensity of calcium (GCaMP6m, E), active Rho (F), and ZO-1 (G) at the site of ZO-1 loss over time in vehicle (water, solid lines) or MSC inhibition (GsMTx4, dashed lines) embryos. Shaded region represents S.E.M. Vehicle: n= 13 flares, 6 embryos, 6 experiments; GsMTx4: n=17 flares, 7 embryos, 6 experiments. (E’-G’) Area under the curve for calcium (E’) and active Rho (F’) were calculated from E and F, respectively. (G’) Scatter plot of change in ZO-1 intensity (G’) was calculated from G. I_1_ and I_2_ represent average intensity of ZO-1 from individual traces at time -25:25 s and 400:450 s, respectively. Error bars represent mean ± S.E.M.; significance calculated using Mann-Whitney test.

### Mechanosensitive calcium channel-mediated calcium influx is required for proper epithelial barrier function and local junction contraction

Rho flare-mediated ZO-1 reinforcement is required to repair local leaks and maintain paracellular barrier integrity. Our results show that a calcium increase follows local leaks (Fig. 1 B), and blocking MSCs leads to impaired Rho activation and ZO-1 reinforcement (Fig. 4, F and G). Therefore, we hypothesized that MSC-mediated calcium influx is required to maintain an intact paracellular epithelial barrier. To detect transient leaks in the barrier, we performed ZnUMBA in vehicle- or GsMTx4-treated embryos. With GsMTx4 treatment, the whole-field intensity of FluoZin-3 increased over time compared to vehicle controls (Fig. 5, A and B), indicating a global barrier disruption. However, global barrier disruption could be the result of failure to repair local leaks associated with ZO-1 loss. Therefore, we next analyzed the frequency of Rho flares in vehicle- and GsMTx4-treated embryos. The frequency of Rho flares increased significantly in GsMTx4-treated embryos compared to the matched controls (Fig. 5 C and Video 7), suggesting an increase in local leaks. To examine if local leaks were contributing to global barrier disruption, we analyzed individual junctions associated with local FluoZin-3 increase. With GsMTx4 treatment, local FluoZin-3 increased repeatedly at the site of ZO-1 loss (single and double white arrows, Fig. 5 E) compared to vehicle controls where the local FluoZin-3 increase was resolved following robust activation of Rho flares and ZO-1 reinforcement (Fig. 5, D-E and Video 8). Of note, in GsMTx4-treated embryos, a Rho flare failed to be activated following an initial increase in local FluoZin-3 (yellow arrow, Fig. 5 E). Collectively, these findings suggest that MSC-mediated robust activation of Rho flares and ZO-1 reinforcement is required to resolve local leaks to maintain the overall paracellular barrier integrity.

**Figure 5:**
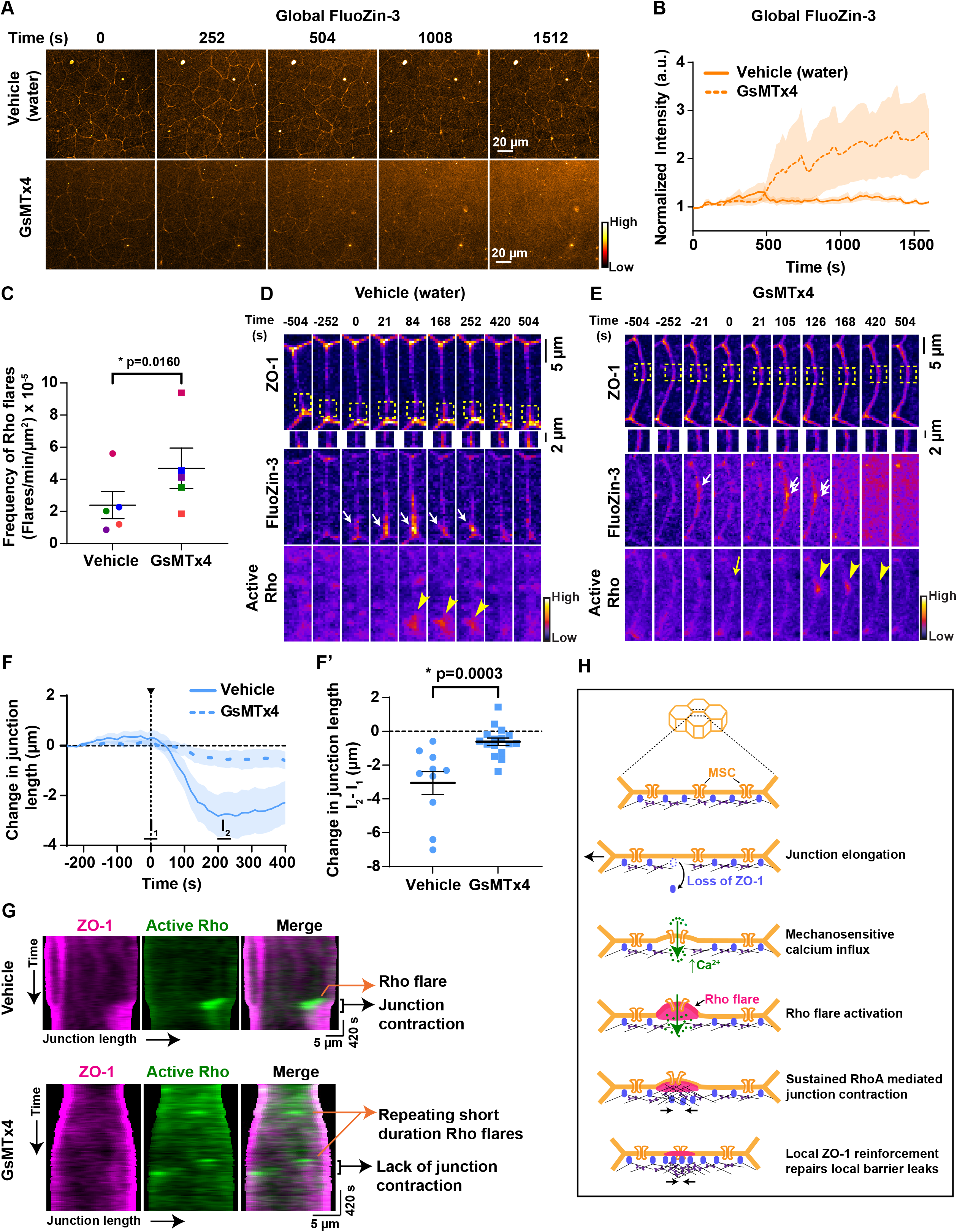
Mechanosensitive calcium channel-mediated calcium influx is required for proper epithelial barrier function and local junction contraction. (A) Time-lapse images (orange hot LUT) of epithelial permeability tested using ZnUMBA (FluoZin-3). Embryos were treated with water (vehicle, top) or 12.5 µM GsMTx4 (MSC inhibitor, bottom). Time 0 s represents the start of the time-lapse movie. (B) Graph of mean normalized intensity of whole-field FluoZin-3 for vehicle (water) or GsMTx4-treated embryos. Blocking MSCs increased global FluoZin-3 intensity over time compared to vehicle. Shaded region represents S.E.M. Vehicle: n= 6 embryos, 4 experiments; GsMTx4: n= 6 embryos, 4 experiments. (C) Frequency of Rho flares in vehicle (water) and GsMTx4-treated embryos. Frequencies from paired experiments are color matched. Error bars represent mean ± S.E.M.; significance calculated using paired two-tailed t-test; n=5 embryos, 4 experiments. (D-E) Montage of representative junctions from A (FIRE LUT). Blocking MSCs with GsMTx4 causes repeated increases in local FluoZin-3 (white arrows), reduced ZO-1 (BFP-ZO-1) reinforcement (yellow boxes, enlarged below), and reduced duration of Rho flares (mCherry-2xrGBD, yellow arrowheads) at sites of ZO-1 loss. Repeated FluoZin-3 increase is indicated by double white arrows, and failure to activate Rho flare after FluoZin-3 increase is indicated by yellow arrow. (F-F’) Graph of change in junction length for junctions with Rho flares in vehicle (water) and GsMTx4-treated embryos. Shaded region represents S.E.M. (F’) Scatter plot of change in junction length was calculated from F. I_1_ and I_2_ represent average length of junction from individual traces from time -20-20 s and 200-240 s, respectively. Error bars represent mean ± S.E.M.; significance calculated using Mann-Whitney test. Vehicle: n= 10 flares, 6 embryos, 6 experiments; GsMTx4: n=16 flares, 7 embryos, 6 experiments. (G) Kymographs of BFP-ZO-1 (magenta) and active Rho (mChe-2xrGBD, green) from representative junctions projected vertex-to-vertex over time for vehicle (water) and GsMTx4-treated embryos. Kymographs highlight the repeating, short duration Rho flares, loss of local junction contraction, and reduced ZO-1 reinforcement upon GsMTx4 treatment. (H) Model showing mechanism by which mechanosensitive calcium signaling regulates epithelial barrier function. Top: 3D view of epithelial cells. Bottom: *en face* view of the junction highlighted showing the spatiotemporal calcium and RhoA signaling following mechanical stimuli.

Following TJ breaks, Myosin II activation through a Rho/ROCK-mediated signaling pathway is required for efficient ZO-1 reinforcement via local junction contraction (Stephenson et al., 2019). So, we asked whether reduced ZO-1 reinforcement observed in GsMTx4-treated embryos was a result of reduced junction contraction. We found that rapid contraction of the junction immediately following the Rho flare was significantly reduced in GsMTx4-treated embryos compared to vehicle controls (Fig. 5, F and F’). Further, to track the local contractile regions within individual junctions, we generated kymographs of a cell-cell junction projected from vertex-to-vertex. In GsMTx4-treated embryos, we observed a dramatic loss of junction contraction and lack of ZO-1 reinforcement following Rho flares (black bracket, Fig. 5 G) compared to robust contraction and ZO-1 reinforcement in controls. Additionally, kymographs highlighted that in GsMTx4-treated embryos there are many short-lived Rho flares, which often repeat at the same site and yet remain unable to reinforce ZO-1 (orange arrows, Fig. 5 G and Fig. S3 A). Furthermore, we observed reduced F-actin accumulation at the site of short-lived Rho flares in GsMTx4-treated embryos (Fig. S3, B-C and Video 9). Taken together, our results show that MSC-mediated calcium influx is required for actomyosin mediated local junction contraction through robust and sustained Rho activation, and thereby efficient reinforcement of ZO-1.

## Discussion

Epithelial tissues adapt to mechanical cues by remodeling their cell-cell junctions and associated cytoskeleton to maintain tissue homeostasis and barrier function. We previously showed that physiological mechanical stimuli – like junction elongation that happens in response to cell shape changes – cause short-lived, local paracellular leaks (Stephenson et al., 2019). However, it was unclear how cells sense local leaks and remodel cell-cell junctions without compromising overall barrier integrity.

Using genetically encoded calcium probes, we observed intracellular calcium flashes during TJ remodeling. Calcium flashes in our study were of longer duration (72.72 ± 9.13 s) compared to single-cell calcium transients (∼26-40 s) that mediate apical cell constriction during *Xenopus* neural tube closure (Christodoulou and Skourides, 2015; Suzuki et al., 2017). The difference in the duration of these calcium transients could be that during apical constriction, the calcium increase is mediated by intracellular calcium stores. Previous studies have shown that intracellular calcium increase precedes TJ formation and establishment of epithelial barrier function (Gonzalez-Mariscal et al., 1990; Nigam et al., 1992). However, these studies used a calcium switch model and found that the timescale of TJ formation following calcium increase is significantly slower (∼50 minutes) than that of TJ reinstatement following a barrier breach and calcium flash described here (∼1-2 minutes). In addition, we demonstrate that local leaks and ZO-1 reduction precede the calcium flash, suggesting that the calcium increase is a consequence of paracellular leak rather than a cause. Thus, our work is the first to suggest that intracellular calcium is a critical regulator of barrier function when cells change shape in an intact epithelium.

Our results also revealed that the intracellular calcium increase originated at the plasma membrane following a leak (Fig. 1, D, and Fig. S1, D-E), suggesting that the calcium increase could be mediated locally by plasma membrane-localized calcium channels or a local membrane rupture. However, previous work from our lab suggested that the plasma membrane is indeed intact at the site of leak (Stephenson et al., 2019). Furthermore, we think it is likely that plasma membrane tension changes locally when the transmembrane TJ proteins uncouple from the cortical actin cytoskeleton upon loss of the scaffolding protein, ZO-1. We demonstrated that calcium increase following local ZO-1 loss is mediated by MSCs, supporting the idea that in epithelial cells, change in local membrane tension led to local activation of MSCs (Shi et al., 2018). Given that both the Piezo and TRP families are mammalian MSCs capable of gating calcium influx, it will be interesting in future studies to investigate the relative role of MSCs blocked by GsMTx4, specifically Piezo1 and TRPC6, in TJ remodeling (Bae et al., 2011; Spassova et al., 2006). Further, it is clear that local calcium increase spreads throughout the cell, thus we cannot rule out the possibility that MSC-mediated calcium influx induces calcium release from intracellular calcium stores.

Both calcium transients and RhoA activation are modulated locally and globally in response to mechanical stimuli (Acharya et al., 2018; Benink and Bement, 2005; Clark et al., 2009; Pardo-Pastor et al., 2018). Our results demonstrate intracellular calcium increase as a key signal for activation of Rho flares. We observed that perturbation of intracellular calcium using a cytoplasmic calcium chelator in conjunction with an IP_3_R blocker (BAPTA-AM+2-APB) reduced RhoA activation at sites of ZO-1 loss to a greater extent compared to the MSC blocker (GsMTx4). The difference in the degree of inhibition could be potentially due to altered tissue tension resulting from physical removal of the vitelline for BAPTA-AM+2-APB but not GsMTx4 experiments, or due to a calcium-independent effect of BAPTA-AM on the actin cytoskeleton (Saoudi et al., 2004). Furthermore, our results show that despite the spreading of calcium throughout the cell, Rho flares are spatially confined to the site of paracellular leaks. This suggests that calcium may regulate RhoA activity by modulating the activity of a specific Rho GEF or Rho GAP, not by direct regulation of RhoA. For example, intracellular calcium increase could activate RhoA via PKCα-mediated activation of a Rho GEF (Holinstat et al., 2003), or CaMKII-mediated crosstalk between active Cdc42 and RhoA (Benink and Bement, 2005; Murakoshi et al., 2011). Interestingly, our observation that in GsMTx4-treated cells, Rho flares were activated but at a significantly lower intensity and shorter duration, suggests an additional calcium-independent mechanism to initiate activation of RhoA.

In addition to regulating the activity of RhoA, MSC-mediated intracellular calcium increase is important for reinforcement of ZO-1, given that both intracellular calcium chelation, and blocking MSCs, inhibited reinforcement of ZO-1. This loss of ZO-1 reinforcement is due to reduced active RhoA-mediated junction contraction, presumably due to the loss of ROCK/Myosin II activation at the site of leaks (Stephenson et al., 2019). Previous studies have demonstrated that chelation of intracellular calcium or inhibition of RhoA activity hinder the association of ZO-1 with the actin cytoskeleton (Nusrat et al., 1995; Stuart et al., 1994). Thus, our study is the first to reveal that calcium promotes ZO-1 recruitment and reinforcement at TJs by regulating RhoA activation. Given that myosin light chain kinase (MLCK) is activated by calcium/calmodulin, and active MLCK stabilizes ZO-1 at TJs (Yu et al., 2010), it is possible that calcium regulates ZO-1 reinforcement at TJs via MLCK activation in parallel to RhoA activation.

Expression and activity of MSCs, including Piezo1 and TRPV4, enhanced global barrier function by up-regulation or post-translational modification of junctional proteins (Akazawa et al., 2013; Sokabe et al., 2010; Zhong et al., 2020). Consistent with this finding, using live imaging and a highly sensitive barrier assay, we observed MSC-mediated calcium influx is required to maintain global barrier function. Interestingly, the increased global paracellular permeability appears to be a cumulative effect of failure to fix local leaks. When MSCs were blocked, Rho flares often failed to be activated immediately following the leak, and when Rho flares were weakly activated, they tended to repeat at the site of the leak in attempts to reinforce ZO-1.

In conclusion, we find mechanically-triggered calcium influx is an important feature of the mechanism by which epithelial cells sense and respond to transient leaks in barrier function (Fig. 5 H). Calcium flashes mediated by MSCs are required for local junction contraction through robust and sustained Rho activation and thereby efficient TJ reinforcement. However, when local leaks fail to be repaired by low intensity, short duration Rho flares, global barrier function is weakened. Thus, MSC-mediated local calcium influx may serve as a mechanotransduction mechanism to specifically relay biochemical signals only to sites of paracellular leaks, without affecting overall TJ structure and function in a dynamic epithelial tissue undergoing various cell shape changes.

## Materials and Methods

### DNA constructs and mRNA preparation

GCaMP6m was generated by PCR amplifying the GCaMP6m sequence from pGP-CMV-GCaMP6m (Addgene plasmid #40754, (Chen et al., 2013)) and ligating it into the *BamH1* and *EcoR1* sites in pCS2+. BFP-PKC-β-C2 and mNeon-PKC-β-C2 were generated by PCR amplifying the C2 domain of *Xenopus laevis* PKC-β from pCS2+/eGFP-PKC-β-C2 (Yu and Bement, 2007) and ligating into the *XhoI* and *EcoR1* sites in pCS2+/N-tagBFP2.0 and pCS2+/N-mNeon, respectively. All constructs were verified by sequencing. pCS2+/mCherry-2xrGBD (probe for active Rho, (Davenport et al., 2016)), pCS2+/BFP-ZO-1 (human ZO-1, (Stephenson et al., 2019)), and pCS2+/BFP-membrane (probe for membrane, (Higashi et al., 2016)) were previously reported. All plasmid DNAs were linearized with *NotI*, except BFP-ZO-1 which was linearized using *KpnI*. mRNA was *in vitro* transcribed from linearized pCS2+ plasmids using the mMessage mMachine SP6 Transcription kit (Ambion) and purified using the RNeasy Mini kit (Qiagen). Transcript size was verified on 1% agarose gel containing 0.05% bleach and Millennium RNA markers (Invitrogen).

### *Xenopus* embryos and microinjection

All studies conducted using *Xenopus laevis* embryos strictly adhered to the compliance standards of the U.S. Department of Health and Human Services Guide for the Care and Use of Laboratory Animals and were approved by the University of Michigan Institutional Animal Care and Use Committee. Eggs from *Xenopus* were collected, *in vitro* fertilized, and dejellied as previously reported (Miller and Bement, 2009; Woolner et al., 2009). Dejellied embryos were stored at 15**°**C in 0.1x Mark’s Modified Ringers solution (MMR) containing 10 mM NaCl, 0.2 mM KCl, 0.2 mM CaCl_2_, 0.1 mM MgCl_2_, and 0.5 mM HEPES, pH 7.4. Embryos were microinjected in the animal hemisphere with mRNA either twice per cell at the two-cell or once per cell at the four-cell stage. Injected embryos were allowed to develop to gastrula stage (Nieuwkoop and Faber stage 10.5–12) at 15**°**C or 17**°**C. The amount of mRNA per 5nl of microinjection volume was as follows: BFP-PKC-β-C2, 100 pg; mNeon-PKC-β-C2, 100 pg; GCaMP6m, 500 pg; mCherry-2xrGBD, 50 pg; BFP-ZO-1, 50 pg; BFP-membrane, 12.5 pg and Lifeact-GFP 10 pg.

### Microscope image acquisition

Live imaging of gastrula stage *Xenopus* embryos was performed on an inverted Olympus Fluoview 1000 Laser Scanning Confocal Microscope equipped with a 60X supercorrected Plan Apo N 60XOSC objective (numerical aperture (NA) = 1.4, working distance = 0.12mm) using mFV10-ASW software. Embryos were imaged in a custom chamber made of a 0.8 mm thick metal slide with a 5 mm hole in the center and two coverslips stuck on both sides of the hole with a thin layer of vacuum grease, lightly compressing the embryo between the coverslips. Embryos were mounted in 0.1x MMR, and the imaging chamber was inverted to image the epithelial cells in the animal hemisphere at room temperature using the Olympus Fluoview 1000 as previously described (Reyes et al., 2014).

Live imaging of calcium and Rho flares was generally captured by scanning the top 3 apical Z-planes (step size of 0.5 µm) and acquired sequentially by line scanning to avoid bleed-through between channels. Global calcium (GCaMP6m), Rho flares, and ZO-1 were acquired at a 2 µs/pixel scanning speed, 5 s time interval, and 1.5-2x zoom. Local calcium (mNeon-PKC-β-C2), Rho flares, and membrane or ZO-1 were acquired at a 4 µs/pixel scanning speed, 10 s time interval, and 2x zoom. For ZnUMBA, 8 apical Z-planes were acquired at an 8 µs/pixel scanning speed, 21 s time interval, and 1.5x zoom.

### Drug treatments

The intracellular calcium chelator BAPTA-AM (Cayman, 15551) and IP_3_R blocker 2-APB (Sigma, D9754) were resuspended in DMSO as a 20 mM or 25 mM stock, respectively, aliquoted and stored at -20°C. Prior to treatment, the translucent vitelline envelope of gastrula-stage embryos was physically removed using sharp forceps as previously described (Sive et al., 2007). Naked embryos were allowed to recover in 0.1x MMR for 30 minutes. Following recovery, embryos were incubated for 1 hour prior to imaging at 15°C in either 0.5% DMSO (vehicle) or a mixture of 20 µM BAPTA-AM and 100 µM 2-APB in 0.1x MMR.

The MSC blocker GsMTx4 (Smartox, 08GSM001) was resuspended in water as a 500 µM stock, aliquoted, and stored at -20°C. Prior to use, GsMTx4 was diluted to a concentration of 25 µM in water, mixed with mRNA, and a total of 1ng was microinjected into each embryo at the 2-4 cell stage.

### Live imaging barrier assay

Zinc-based Ultrasensitive Microscopic Barrier Assay (ZnUMBA) was performed as previously reported (Stephenson et al., 2019). Gastrula-stage albino embryos expressing mCherry-2xrGBD and BFP-ZO-1 or BFP-PKC-β-C2 were microinjected with 10 nl of 1 mM FluoZin-3 containing 100 µM CaCl_2_ and 100 µM EDTA into the blastocoel. FluoZin-3-injected embryos were allowed to recover from microinjection for 5 minutes in 0.1x MMR. Following recovery, embryos were mounted in 0.1x MMR containing 1 mM ZnCl_2_ and imaged immediately using confocal microscopy.

### Immunofluorescence staining

Albino embryos injected with vehicle (water) or 12.5 µM GsMTx4 were fixed and immunostained as previously reported (Arnold et al., 2019). At gastrula-stage, embryos were fixed overnight at room temperature with a mix of 1.5% PFA, 0.25% glutaraldehyde, 0.2% Triton X-100, Alexa Fluor 647 phalloidin (1:1000, Thermo Fisher Scientific, # A22287) in a 0.88x MT buffer containing 80 mM K-PIPES, 5 mM EGTA, and 1 mM MgCl_2_, pH 6.8 with KOH. Fixed embryos were quenched for 1 hour at room temperature with 100mM sodium borohydride in 1x PBS and blocked overnight in 10% FBS, 5% DMSO and 0.1% NP-40 in 1x Tris-buffered Saline. Animal hemisphere of the bisected embryos were incubated overnight in anti-P-MLC (1:100, Cell Signaling Technologies, # 3671) in blocking solution, washed 3x with blocking solution, and incubated overnight in anti-rabbit Alexa 488 IgG (1:200, Thermo Fisher Scientific, # A11008) and Alexa Fluor 647 phalloidin (1:1000). Embryos were washed and mounted in VECTASHIELD (VWR, # 101098-042).

### Image processing and quantification

Image processing and analysis were performed using Image J (Fiji). Sum intensity projections of the Z-series were used for all quantification, and confocal images of time-lapse movies represented in all figures are sum projections of 3 apical Z-planes (1.5 µm), with the exception of FluoZin-3 (8 apical Z-planes, 4 µm).

### Quantification of calcium dynamics during Rho flares

Calcium dynamics at Rho flares were measured using a circular region of interest of 2.5 µm diameter drawn at the site of flares, spanning the membrane and cytoplasm adjacent to the membrane, manually tracked over every frame to account for in-plane drifting in live tissue, and normalized to baseline of 1. Cytosolic calcium (GCaMP6m) was calculated for each frame using the formula, F_normalized_ = (F/F_baseline_), where F_baseline_ represents the average calcium intensity of the first 150 s. Local calcium (PKC-C2) was calculated for each frame using the formula, F = (F_flare_ – F_background_) and normalized to a baseline of 1, where F_baseline_ represents the average calcium intensity of first 100 s and F_background_ represents the cytoplasmic C2 intensity measured using an ROI of same size. An average of three independent measurements of calcium dynamics at each Rho flare was considered one data point (n=1).

For accurate measurement of calcium flash dynamics using the cytoplasmic soluble calcium probe (GCaMP6m), Rho flares were selected from cells that exhibited an isolated Rho flare that was not interrupted by a multicellular travelling calcium wave for 500 s prior to and after the initiation of Rho flares.

Intensity and duration of Rho flares and ZO-1 were measured at the site of Rho flares using a small circular ROI (GCaMP6m: 0.8 µm; PKC-C2: 2.5 µm; and Fluozin-3: 2.2 µm) and normalized to a nearby reference junction without Rho flares using ROI of same size to account for photobleaching and drifting in the Z-plane. The increase in Rho activity at the start of the Rho flare was aligned to time 0 s as previously described (Stephenson et al., 2019), with the exception of Fig. 3. For Fig. 3, as calcium chelation (BAPTA-AM + 2-APB) significantly reduced the intensity of Rho flares, we adopted an alternate method to align time 0 s in the graph to the frame before the rapid decrease in ZO-1; the decrease in ZO-1 was defined as a 5% decrease in normalized ZO-1 intensity for at least 3 out of 4 consecutive frames.

### Duration of calcium flash

Duration of the calcium flash was quantified by measuring the full duration at half-maximum (FDHM) of the individual normalized calcium flashes at sites of Rho flares. FDHM (seconds) was measured manually by calculating the duration at 50% of the maximum amplitude above the baseline of 1 on the ascent and descent of the calcium transient. Experiments where calcium increased less than two-fold over the baseline were eliminated from the analysis. FDHM was plotted in a scatter plot.

### Area under the curve

Area under the curve for active Rho flares and calcium flashes was quantified using GraphPad Prism 9.0 by applying area under the curve (AUC) analysis of the individual traces. AUC analysis was applied to all points in the peaks above the baseline of 1, and peaks less than 10% of the increase from the baseline to the maximum Y-value were ignored. Individual AUC values for each Rho flare or calcium flash were plotted in a scatter plot.

### Frequency of Rho flares

Rho flare frequency was measured manually using Image J, by counting the number of Rho flare occurrences at cell-cell junctions in the whole field of imaging from the start of the time-lapse movie. Rho flares where the local active RhoA intensity increased and was sustained at the junctions for a span of 4-5 frames (84 s-105 s) was included in the analysis.

### Global barrier function

Global barrier function was quantified by measuring the whole field intensity of Fluozin-3 signal, including junctional and cytoplasmic, over time using an ROI of 141.12×141.12 µm. Time 0 s represents the start of image acquisition, which began immediately following mounting of the embryo injected with Fluozin-3 in 0.1x MMR containing 1 mM ZnCl_2_. Each timepoint was measured, and baseline was normalized to 1 by dividing the individual value by the average of the first 84 seconds.

### Junction length

Length of the cell-cell junction was quantified using ZO-1 as junctional marker. Using Image J, a 0.3 µm wide segmented line was drawn on the junction with Rho flares to trace the junction from vertex-to-vertex. Length of the line was measured for every other frame by manually advancing and adjusting the line to accommodate cell shape changes through the time lapse movie. Each junction was measured in triplicate, and baseline was normalized to 1 by dividing the individual value by the average of the first 6 data points.

### Construction of kymographs

Kymographs from vertex-to-vertex of a cell-cell junction were constructed as previously described (Stephenson et al., 2019). Briefly, cell-cell junctions were digitized using ZO-1 as junctional marker. Each horizontal line in the kymograph was generated by measuring the intensity of ZO-1 and active Rho using 0.75 µm circular ROIs positioned at points along the length of the cell-cell junction. Kymographs were constructed by stacking and center aligning the horizontal lines from successive time frames.

### Apical cell perimeter

Apical perimeter of the cell was quantified using ZO-1 to identify cell boundaries. Using Image J, a segmented line was drawn to trace the boundary of the cells with a calcium flash and neighboring cells without a calcium flash over time. The perimeter of the cell was measured for every time point by manually adjusting the line to accommodate cell shape changes. Change in apical perimeter was calculated by subtracting the current perimeter from the initial perimeter (average perimeter over the first 3 time points).

### Intensity of F-actin and P-MLC

Junctional and cytoplasmic intensity of F-actin and P-MLC were quantified in Image J. Junctional intensity was measured using a 2.07 µm wide segmented line to trace a bicellular junction (excluding vertices) and the matched cytoplasmic intensity was measured using a 10.4 µm circular ROI. Ratio of junctional/cytoplasmic F-actin or P-MLC was calculated by dividing the intensity at the junction by the intensity in the cytoplasm for each cell.

### Statistical analysis

Statistical analysis and standard error of the mean (S.E.M.) were calculated in GraphPad Prism 9.0. Statistical analysis between two groups was measured using a two-tailed, unpaired Mann-Whitney U test, with the exception of flare frequency and intensity of actomyosin. The statistical significance of frequency of Rho flares was measured using a two-tailed, paired Student’s *t*-test. Statistical significance of F-actin and P-MLC intensity at the junctions and junction/cytoplasm ratio was measured using a One-way ANOVA test.

## Acknowledgements

We thank W.M.Bement for pCS2+/ mCherry-2xrGBD and pCS2+/ eGFP-PKC-β-C2 constructs as well as current and former members of the A.L.Miller laboratory for providing helpful discussion and feedback on experiments. This work was supported by NIH grant (2R01 GM112794) to A.L.Miller, a predoctoral fellowship from the American Heart Association (20PRE35120588) to S.Varadarajan, and BBSRC grants BB/P01190X and BB/P006507 to A.B.Goryachev.

The authors declare no competing financial interests.

## Author contributions

S.Varadarajan and A.L.Miller conceptualized the study; S.Varadarajan, R.E.Stephenson, and A.L.Miller developed the methodology; S.Varadarajan performed the majority of experiments and data analysis; J.L.Wu performed the experiment and analyzed data for Fig. S3; E.R.Misterovich analyzed the data for Fig. 2F, Fig. 5 F and F’; I.S.Erofeev constructed the kymographs for Fig. 5G; S.Varadarajan and A.L.Miller wrote the original draft of the manuscript with input from R.E.Stephenson; All authors revised the manuscript; A.L.Miller and A.B.Goryachev acquired funding; A.L.Miller supervised the study.

## Abbreviations

GBD: GTPase-binding domain
GEF: guanine nucleotide exchange factor
GAP: GTPase-activating protein
GsMTx4: Grammostola Mechanotoxin #4
LUT: lookup table
MSC: mechanosensitive calcium channel
RhoA-GTP: active RhoA
ROI: region of interest
TRP: Transient Receptor Potential
ZnUMBA: Zinc-based Ultrasensitive Microscopic Barrier Assay

## References

Acharya, B.R., A. Nestor-Bergmann, X. Liang, S. Gupta, K. Duszyc, E. Gauquelin, G.A. Gomez, S. Budnar, P. Marcq, O.E. Jensen, Z. Bryant, and A.S. Yap. 2018. A Mechanosensitive RhoA Pathway that Protects Epithelia against Acute Tensile Stress. Developmental cell. 47:439–452 e436.

Akazawa, Y., T. Yuki, H. Yoshida, Y. Sugiyama, and S. Inoue. 2013. Activation of TRPV4 strengthens the tight-junction barrier in human epidermal keratinocytes. Skin Pharmacol Physiol. 26:15–21.

Arnold, T.R., J.H. Shawky, R.E. Stephenson, K.M. Dinshaw, T. Higashi, F. Huq, L.A. Davidson, and A.L. Miller. 2019. Anillin regulates epithelial cell mechanics by structuring the medial-apical actomyosin network. Elife. 8.

Bae, C., F. Sachs, and P.A. Gottlieb. 2011. The mechanosensitive ion channel Piezo1 is inhibited by the peptide GsMTx4. Biochemistry. 50:6295–6300.

Benink, H.A., and W.M. Bement. 2005. Concentric zones of active RhoA and Cdc42 around single cell wounds. Journal of Cell Biology. 168:429–439.

Bowman, C.L., P.A. Gottlieb, T.M. Suchyna, Y.K. Murphy, and F. Sachs. 2007. Mechanosensitive ion channels and the peptide inhibitor GsMTx-4: history, properties, mechanisms and pharmacology. Toxicon. 49:249–270.

Canales, J., D. Morales, C. Blanco, J. Rivas, N. Diaz, I. Angelopoulos, and O. Cerda. 2019. A TR(i)P to Cell Migration: New Roles of TRP Channels in Mechanotransduction and Cancer. Front Physiol. 10:757.

Cavanaugh, K.J., T.S. Cohen, and S.S. Margulies. 2006. Stretch increases alveolar epithelial permeability to uncharged micromolecules. American journal of physiology. Cell physiology. 290:C1179–1188.

Chen, T.W., T.J. Wardill, Y. Sun, S.R. Pulver, S.L. Renninger, A. Baohan, E.R. Schreiter, R.A. Kerr, M.B. Orger, V. Jayaraman, L.L. Looger, K. Svoboda, and D.S. Kim. 2013. Ultrasensitive fluorescent proteins for imaging neuronal activity. Nature. 499:295–300.

Christodoulou, N., and P.A. Skourides. 2015. Cell-Autonomous Ca(2+) Flashes Elicit Pulsed Contractions of an Apical Actin Network to Drive Apical Constriction during Neural Tube Closure. Cell reports. 13:2189–2202.

Clark, A.G., A.L. Miller, E. Vaughan, H.Y.E. Yu, R. Penkert, and W.M. Bement. 2009. Integration of Single and Multicellular Wound Responses. Current Biology. 19:1389–1395.

Claude, P., and D.A. Goodenough. 1973. Fracture faces of zonulae occludentes from “tight” and “leaky” epithelia. The Journal of cell biology. 58:390–400.

Cohen, T.S., K.J. Cavanaugh, and S.S. Margulies. 2008. Frequency and peak stretch magnitude affect alveolar epithelial permeability. Eur Respir J. 32:854–861.

Coste, B., J. Mathur, M. Schmidt, T.J. Earley, S. Ranade, M.J. Petrus, A.E. Dubin, and A. Patapoutian. 2010. Piezo1 and Piezo2 Are Essential Components of Distinct Mechanically Activated Cation Channels. Science. 330:55–60.

Davenport, N.R., K.J. Sonnemann, K.W. Eliceiri, and W.M. Bement. 2016. Membrane dynamics during cellular wound repair. Molecular biology of the cell. 27:2272–2285.

Ellefsen, K.L., J.R. Holt, A.C. Chang, J.L. Nourse, J. Arulmoli, A.H. Mekhdjian, H. Abuwarda, F. Tombola, L.A. Flanagan, A.R. Dunn, I. Parker, and M.M. Pathak. 2019. Myosin-II mediated traction forces evoke localized Piezo1-dependent Ca(2+) flickers. Commun Biol. 2:298.

Fanning, A.S., B.J. Jameson, L.A. Jesaitis, and J.M. Anderson. 1998. The tight junction protein ZO-1 establishes a link between the transmembrane protein occludin and the actin cytoskeleton. J Biol Chem. 273:29745–29753.

Furuse, M., H. Sasaki, K. Fujimoto, and S. Tsukita. 1998. A single gene product, claudin-1 or -2, reconstitutes tight junction strands and recruits occludin in fibroblasts. The Journal of cell biology. 143:391–401.

Gnanasambandam, R., C. Ghatak, A. Yasmann, K. Nishizawa, F. Sachs, A.S. Ladokhin, S.I. Sukharev, and T.M. Suchyna. 2017. GsMTx4: Mechanism of Inhibiting Mechanosensitive Ion Channels. Biophysical Journal. 112:31–45.

Gonzalez-Mariscal, L., R.G. Contreras, J.J. Bolivar, A. Ponce, B. Chavez De Ramirez, and M. Cereijido. 1990. Role of calcium in tight junction formation between epithelial cells. The American journal of physiology. 259:C978–986.

Gudipaty, S.A., J. Lindblom, P.D. Loftus, M.J. Redd, K. Edes, C.F. Davey, V. Krishnegowda, and J. Rosenblatt. 2017. Mechanical stretch triggers rapid epithelial cell division through Piezo1. Nature. 543:118–121.

Gudipaty, S.A., and J. Rosenblatt. 2016. Epithelial cell extrusion: Pathways and pathologies. Seminars in Cell and Developmental Biology.

Guillot, C., and T. Lecuit. 2013. Mechanics of epithelial tissue homeostasis and morphogenesis. Science. 340:1185–1189.

Higashi, T., T.R. Arnold, R.E. Stephenson, K.M. Dinshaw, and A.L. Miller. 2016. Maintenance of the Epithelial Barrier and Remodeling of Cell-Cell Junctions during Cytokinesis. Current biology : CB. 26:1829–1842.

Holinstat, M., D. Mehta, T. Kozasa, R.D. Minshall, and A.B. Malik. 2003. Protein kinase Calpha-induced p115RhoGEF phosphorylation signals endothelial cytoskeletal rearrangement. The Journal of biological chemistry. 278:28793–28798.

Hunter, G.L., J.M. Crawford, J.Z. Genkins, and D.P. Kiehart. 2014. Ion channels contribute to the regulation of cell sheet forces during Drosophila dorsal closure. Development. 141:325–334.

Itoh, M., M. Furuse, K. Morita, K. Kubota, M. Saitou, and S. Tsukita. 1999. Direct binding of three tight junction-associated MAGUKs, ZO-1, ZO-2, and ZO-3, with the COOH termini of claudins. The Journal of cell biology. 147:1351–1363.

Ivanov, A.I., C.A. Parkos, and A. Nusrat. 2010. Cytoskeletal Regulation of Epithelial Barrier Function During Inflammation. The American journal of pathology. 177:512–524.

Liu, C., and C. Montell. 2015. Forcing open TRP channels: Mechanical gating as a unifying activation mechanism. Biochem Biophys Res Commun. 460:22–25.

Luissint, A.C., C.A. Parkos, and A. Nusrat. 2016. Inflammation and the Intestinal Barrier: Leukocyte-Epithelial Cell Interactions, Cell Junction Remodeling, and Mucosal Repair. Gastroenterology. 151:616–632.

Marchiando, A.M., W.V. Graham, and J.R. Turner. 2010. Epithelial barriers in homeostasis and disease. Annual review of pathology. 5:119–144.

Miller, A.L., and W.M. Bement. 2009. Regulation of cytokinesis by Rho GTPase flux. Nat Cell Biol. 11:71–77.

Miyamoto, T., T. Mochizuki, H. Nakagomi, S. Kira, M. Watanabe, Y. Takayama, Y. Suzuki, S. Koizumi, M. Takeda, and M. Tominaga. 2014. Functional role for Piezo1 in stretch-evoked Ca2+ influx and ATP release in Urothelial cell cultures. Journal of Biological Chemistry. 289:16565–16575.

Mochizuki, T., T. Sokabe, I. Araki, K. Fujishita, K. Shibasaki, K. Uchida, K. Naruse, S. Koizumi, M. Takeda, and M. Tominaga. 2009. The TRPV4 cation channel mediates stretch-evoked Ca2+ influx and ATP release in primary urothelial cell cultures. The Journal of biological chemistry. 284:21257–21264.

Moriwaki, K., S. Tsukita, and M. Furuse. 2007. Tight junctions containing claudin 4 and 6 are essential for blastocyst formation in preimplantation mouse embryos. Dev Biol. 312:509–522.

Murakoshi, H., H. Wang, and R. Yasuda. 2011. Local, persistent activation of Rho GTPases during plasticity of single dendritic spines. Nature. 472:100–104.

Nigam, S.K., E. Rodriguez-Boulan, and R.B. Silver. 1992. Changes in intracellular calcium during the development of epithelial polarity and junctions. Proceedings of the National Academy of Sciences of the United States of America. 89:6162–6166.

Nusrat, A., M. Giry, J.R. Turner, S.P. Colgan, C.A. Parkos, D. Carnes, E. Lemichez, P. Boquet, and J.L. Madara. 1995. Rho protein regulates tight junctions and perijunctional actin organization in polarized epithelia. Proceedings of the National Academy of Sciences of the United States of America. 92:10629–10633.

Pardo-Pastor, C., F. Rubio-Moscardo, M. Vogel-González, S.A. Serra, A. Afthinos, S. Mrkonjic, O. Destaing, J.F. Abenza, J.M. Fernández-Fernández, X. Trepat, C. Albiges-Rizo, K. Konstantopoulos, and M.A. Valverde. 2018. Piezo2 channel regulates RhoA and actin cytoskeleton to promote cell mechanobiological responses. Proceedings of the National Academy of Sciences. 115:1925–1930.

Reyes, C.C., M. Jin, E.B. Breznau, R. Espino, R. Delgado-Gonzalo, A.B. Goryachev, and A.L. Miller. 2014. Anillin regulates cell-cell junction integrity by organizing junctional accumulation of Rho-GTP and actomyosin. Current Biology. 24:1263–1270.

Rosenblatt, J., M.C. Raff, and L.P. Cramer. 2001. An epithelial cell destined for apoptosis signals its neighbors to extrude it by an actin- and myosin-dependent mechanism. Current biology : CB. 11:1847–1857.

Sahu, S.U., M.R. Visetsouk, R.J. Garde, L. Hennes, C. Kwas, and J.H. Gutzman. 2017. Calcium signals drive cell shape changes during zebrafish midbrain–hindbrain boundary formation. Molecular biology of the cell. 28:875–882.

Samak, G., T. Suzuki, A. Bhargava, and R.K. Rao. 2010. c-Jun NH2-terminal kinase-2 mediates osmotic stress-induced tight junction disruption in the intestinal epithelium. American journal of physiology. Gastrointestinal and liver physiology. 299:G572–584.

Saoudi, Y., B. Rousseau, J. Doussiere, S. Charrasse, C. Gauthier-Rouviere, N. Morin, C. Sautet-Laugier, E. Denarier, R. Scaife, C. Mioskowski, and D. Job. 2004. Calcium-independent cytoskeleton disassembly induced by BAPTA. Eur J Biochem. 271:3255–3264.

Shao, X., Q. Li, A. Mogilner, A.D. Bershadsky, and G.V. Shivashankar. 2015. Mechanical stimulation induces formin-dependent assembly of a perinuclear actin rim. Proceedings of the National Academy of Sciences of the United States of America. 112:E2595–2601.

Shi, Z., Z.T. Graber, T. Baumgart, H.A. Stone, and A.E. Cohen. 2018. Cell Membranes Resist Flow. Cell. 175:1769–1779 e1713.

Sive, H.L., R.M. Grainger, and R.M. Harland. 2007. Removing the Vitelline Membrane from Xenopus laevis Embryos. CSH Protoc. 2007:pdb prot4732.

Sokabe, T., T. Fukumi-Tominaga, S. Yonemura, A. Mizuno, and M. Tominaga. 2010. The TRPV4 channel contributes to intercellular junction formation in keratinocytes. The Journal of biological chemistry. 285:18749–18758.

Spassova, M.A., T. Hewavitharana, W. Xu, J. Soboloff, and D.L. Gill. 2006. A common mechanism underlies stretch activation and receptor activation of TRPC6 channels. Proceedings of the National Academy of Sciences. 103:16586–16591.

Staehelin, L.A., T.M. Mukherjee, and A.W. Williams. 1969. Freeze-etch appearance of the tight junctions in the epithelium of small and large intestine of mice. Protoplasma. 67:165–184.

Stephenson, R.E., T. Higashi, I.S. Erofeev, T.R. Arnold, M. Leda, A.B. Goryachev, and A.L. Miller. 2019. Rho Flares Repair Local Tight Junction Leaks. Developmental cell. 48:445–459 e445.

Stuart, R.O., A. Sun, M. Panichas, S.C. Hebert, B.M. Brenner, and S.K. Nigam. 1994. Critical role for intracellular calcium in tight junction biogenesis. J Cell Physiol. 159:423–433.

Suzuki, M., M. Sato, H. Koyama, Y. Hara, K. Hayashi, N. Yasue, H. Imamura, T. Fujimori, T. Nagai, R.E. Campbell, and N. Ueno. 2017. Distinct intracellular Ca(2+) dynamics regulate apical constriction and differentially contribute to neural tube closure. Development. 144:1307–1316.

Van Itallie, C.M., A.J. Tietgens, and J.M. Anderson. 2017. Visualizing the dynamic coupling of claudin strands to the actin cytoskeleton through ZO-1. Mol Biol Cell. 28:524–534.

Varadarajan, S., R.E. Stephenson, and A.L. Miller. 2019. Multiscale dynamics of tight junction remodeling. Journal of cell science. 132.

Wallingford, J.B., A.J. Ewald, R.M. Harland, and S.E. Fraser. 2001. Calcium signaling during convergent extension in Xenopus. Current Biology. 11:652–661.

Wei, C., X. Wang, M. Chen, K. Ouyang, L.S. Song, and H. Cheng. 2009. Calcium flickers steer cell migration. Nature. 457:901–905.

Woolner, S., A.L. Miller, and W.M. Bement. 2009. Imaging the cytoskeleton in live Xenopus laevis embryos. Methods Mol Biol. 586:23–39.

Yu, D., A.M. Marchiando, C.R. Weber, D.R. Raleigh, Y. Wang, L. Shen, and J.R. Turner. 2010. MLCK-dependent exchange and actin binding region-dependent anchoring of ZO-1 regulate tight junction barrier function. Proceedings of the National Academy of Sciences of the United States of America. 107:8237–8241.

Yu, H.Y.E., and W.M. Bement. 2007. Control of local actin assembly by membrane fusion-dependent compartment mixing. Nature Cell Biology. 9:149–159.

Zhong, M., W. Wu, H. Kang, Z. Hong, S. Xiong, X. Gao, J. Rehman, Y.A. Komarova, and A.B. Malik. 2020. Alveolar Stretch Activation of Endothelial Piezo1 Protects Adherens Junctions and Lung Vascular Barrier. Am J Respir Cell Mol Biol. 62:168–177.

Zihni, C., C. Mills, K. Matter, and M.S. Balda. 2016. Tight junctions: From simple barriers to multifunctional molecular gates. Nature Reviews Molecular Cell Biology. 17:564–580.

