## Supplemental material for "Mechanosensitive calcium signaling in response to cell shape changes promotes epithelial tight junction remodeling by activating RhoA"

Figure S1 (related to Figure 1 and 2)

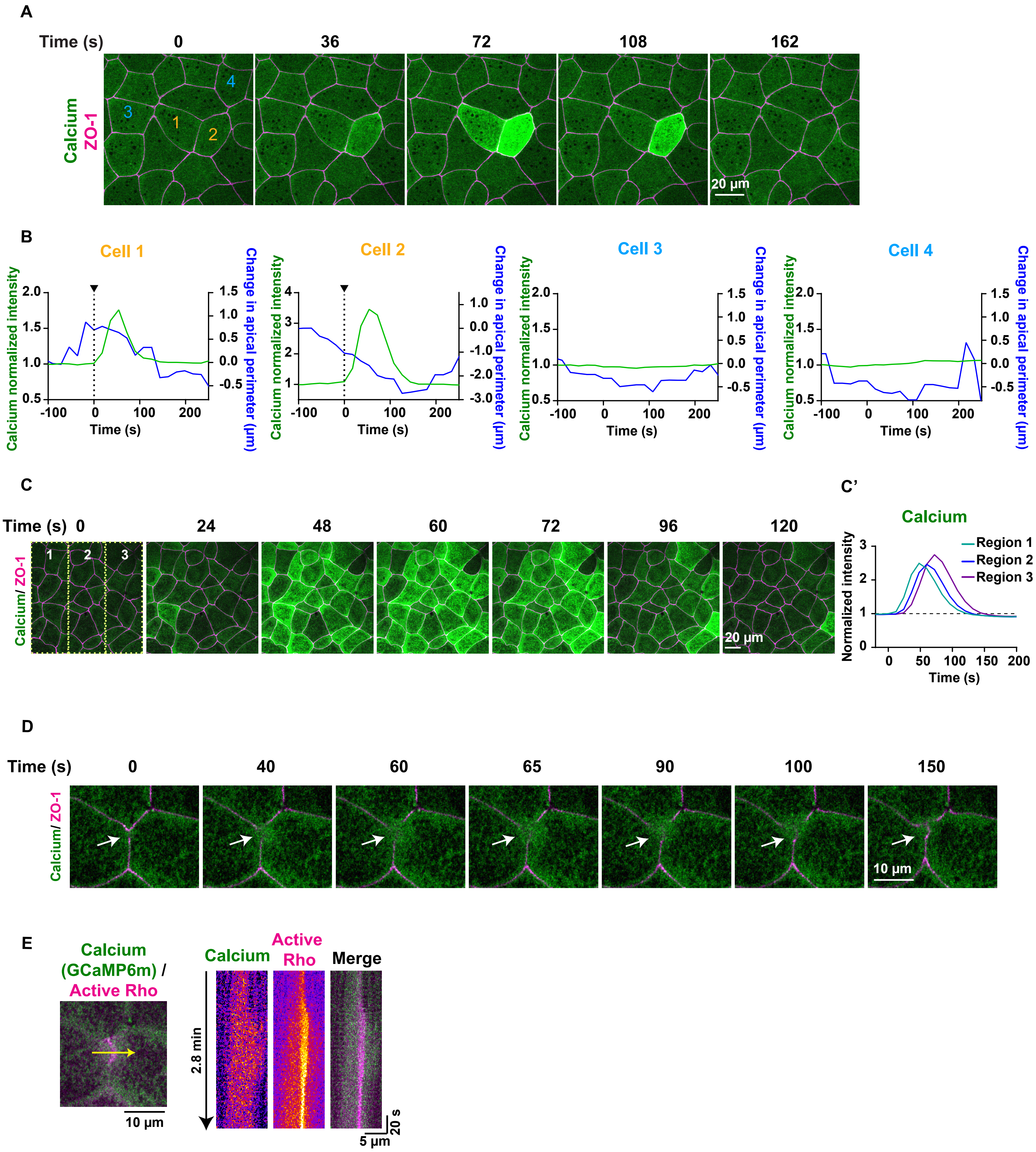

Figure S2 (related to Figure 3 and 4)

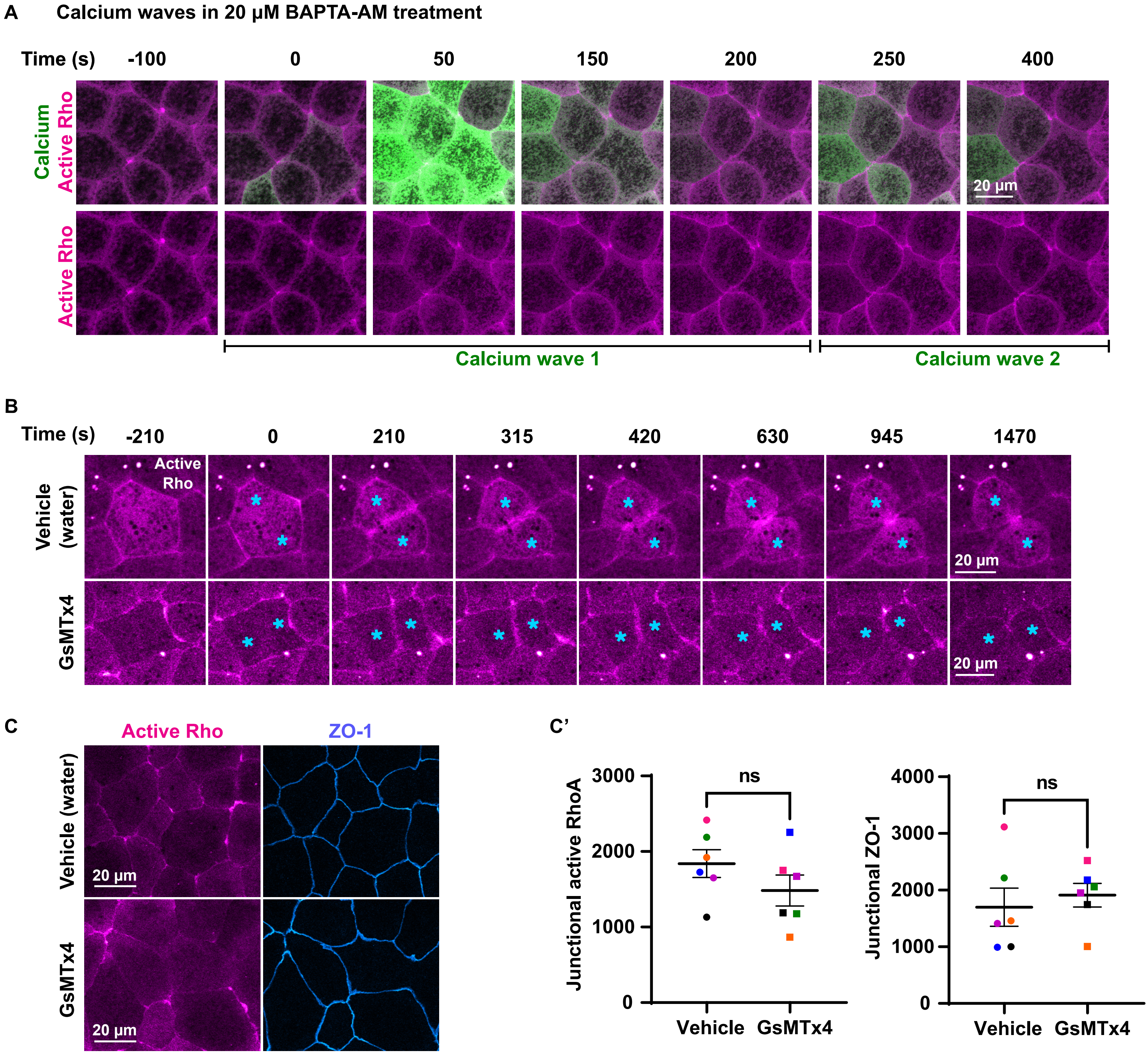

Figure S3 (related to Figure 5)

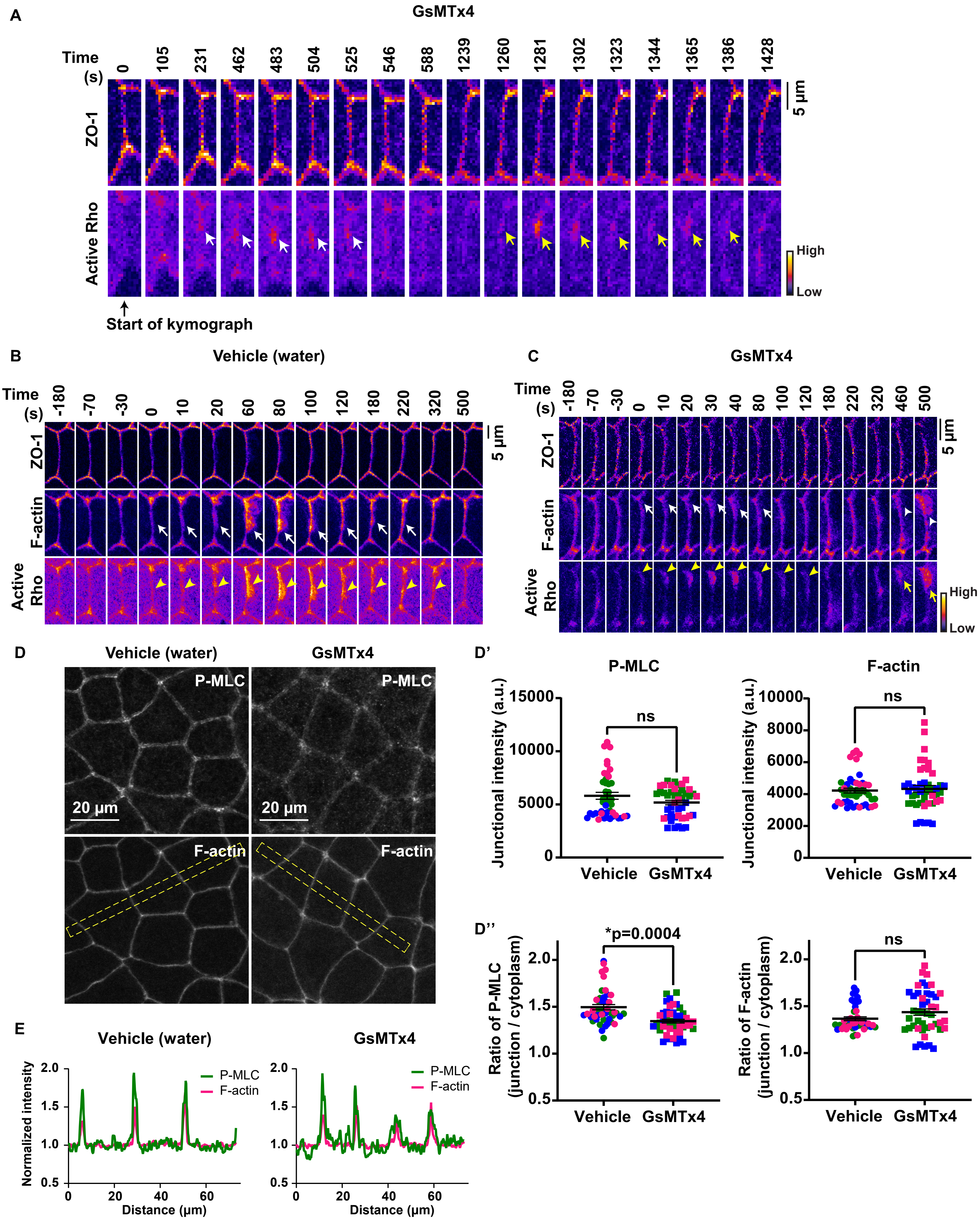

**Supplemental figure legends:**

**Figure S1: Dynamics of calcium flash and calcium waves in gastrula-stage *Xenopus laevis* epithelium**.

(A) Live imaging of calcium (GCaMP6m, green) and ZO-1 (BFP-ZO-1, magenta) in the animal cap epithelium of gastrula-stage *Xenopus* embryos. Calcium flash is short-lived and restricted to cell 1 and cell 2 (orange numbers). Time 0 s represents the start of the calcium flash.

(B) Graphs showing change in cellular apical perimeter and calcium intensity in 4 different cells shown in A over time. Cells experiencing calcium flash (cells 1 and 2) decrease in apical perimeter compared to control cells (cells 3 and 4).

(C-C') Live imaging of calcium (GCaMP6m, green) and ZO-1 (BFP-ZO-1, magenta) in the animal cap epithelium of gastrula-stage *Xenopus* embryos. Travelling calcium wave starts at time 0 s in region 1, propagates through regions 2 and 3, and subsides in ~120 s. (C') Graph showing the increase in calcium intensity over time in regions marked in the first frame of C.

(D) Live imaging of calcium (GCaMP6m, green) and ZO-1 (BFP-ZO-1, magenta) showing a local calcium increase (white arrows) in cells expressing a reduced amount of GCaMP6m. Time 0 s represents the start of the calcium increase.

(E) Left: Cell view of embryo expressing calcium probe (GCaMP6m, green) and active Rho probe (mCherry-2xrGBD, magenta). 5-pixel wide yellow arrow indicates the region used to generate the kymograph. Right: Kymograph shows that cytosolic calcium increases locally at the site of the Rho flare. Individual images are shown using FIRE LUT.

**Figure S2: Mechanosensitive calcium channel-mediated calcium influx is not required for baseline junctional Rho activity or Rho activation at the contractile ring**.

(A) Live imaging of calcium (GCaMP6m, green) and active Rho probe (mCherry-2xrGBD, magenta) in the animal cap epithelium of gastrula-stage *Xenopus* embryos treated with 20 µM BAPTA-AM in DMSO after vitelline removal. Montage shows recurring calcium waves within a span of 500 s. First calcium wave starts at time 0 s, and second calcium wave starts at ~200 s.

(B) Live imaging of active Rho (mCherry-2xrGBD, magenta) in embryos treated with vehicle (water) or 12.5 µM GsMTx4 (MSC inhibitor). Montage shows MSC inhibition does not affect Rho activity at the contractile ring, and cells complete cytokinesis successfully. Blue asterisk indicates dividing cells. Time 0 s represents the start of contractile ring formation.

(C-C') Live imaging of ZO-1 (BFP-ZO-1, blue) and active Rho (mCherry-2xrGBD, magenta) in embryos treated with vehicle (water) or 12.5 μM GsMTx4. (C') Quantification shows that MSC inhibition does not significantly affect baseline junctional RhoA activity at apical cell-cell junctions. Junctional intensity of active Rho and ZO-1 from paired experiments are color matched. Error bars represent mean ± S.E.M.; significance calculated using Wilcoxon matched-pairs test; n=30 junctions, 6 experiments.

**Figure S3: Sustained Rho flares are required for robust F-actin accumulation and successful reinforcement of ZO-1**

(A) Montage of a representative junction used to construct kymograph in Fig. 5G (FIRE LUT). Blocking MSCs with 12.5 μM GsMTx4 causes repeated increases in active Rho at the site of reduced ZO-1 at time 231 s (first flare, white arrows) and 1260 s (second flare, yellow arrows). Note that ZO-1 is partially reinforced following the first flare but breaks at the same site prior to activation of second flare. Time 0 s represents the start of kymograph in Fig. 5G.

(B-C) Time-lapse images (FIRE LUT) of ZO-1 (BFP-ZO-1), F-actin (Lifeact-GFP), and active Rho (mCherry-2xrGBD) in embryos treated with vehicle (water) or 12.5 μM GsMTx4. Blocking MSCs with 12.5 μM GsMTx4 causes reduced F-actin accumulation (C, white arrows) at the site of Rho flares (C, yellow arrowheads) compared to vehicle control in B. Note the robust F-actin accumulation (C, white arrowhead) scales with a higher intensity repeating Rho flare (C, yellow arrow). Time 0 s represents the start of Rho flare.

(D-D'') Sum projection of fixed staining for P-MLC (anti-pMLC) and F-actin (Alexa Fluor 647 phalloidin) in embryos treated with vehicle (water) or 12.5 μM GsMTx4. (D') Quantification shows that MSC inhibition does not significantly affect the junctional intensity of P-MLC and F-actin at cell-cell junctions compared to vehicle control. (D'') MSC inhibition reduces the junction/cytoplasm ratio of P-MLC compared to vehicle, but does not affect the junction/cytoplasm ratio for F-actin. Data points from paired experiments are color matched. Error bars represent mean ± S.E.M.; significance calculated using One-way ANOVA test; n=45 junctions, 3 experiments.

(E) Line scan of P-MLC (green) and F-actin (magenta) across multiple junctions for representative image shown in D (yellow boxes) shows that intensity and width of P-MLC and F-actin peaks at cell-cell junctions in GsMTx4-treated embryos are comparable to vehicle controls.

**Video legends:**

**Video 1:** Time-lapse confocal imaging of gastrula-stage *Xenopus laevis* epithelium showing local calcium (BFP-C2, green) and active Rho (mCherry-2xrGBD, magenta) increase at the site of leaks (FluoZin-3, FIRE LUT). Video illustrates that the FluoZin-3 increase precedes the local calcium increase. Time interval, 21 s; video frame rate, 5 fps.

**Video 2:** Time-lapse confocal imaging shows calcium (mNeon-C2, green) increases locally at sites of Rho flares (mCherry-2xrGBD, magenta) during cell shape changes. Time interval, 10 s; video frame rate, 20 fps. Related to Fig. 1C.

**Video 3:** Time-lapse confocal imaging shows naturally occurring calcium flash and calcium wave in a gastrula-stage *Xenopus* embryo expressing BFP-ZO-1 (magenta) and GCaMP6m (green). Video shows the calcium flash is restricted to 2-3 cells, whereas the calcium wave travels across the tissue. Time interval, 18 s for calcium flash and 12 s for calcium wave; video frame rate, 2 fps. Related to Fig. S1 A-D.

**Video 4:** Time-lapse confocal imaging shows the dynamic activation of Rho flares (mCherry-2xrGBD, magenta) at the site of ZO-1 loss (BFP-ZO-1, cyan) and calcium flashes (GCaMP6m, green). Time interval, 5 s; video frame rate, 20 fps.

**Video 5:** Time-lapse confocal imaging shows higher amplitude (compared with Video 4) calcium flash (GCaMP6m, green) and higher intensity Rho flare (mCherry-2xrGBD, magenta) during dramatic loss and reinforcement of ZO-1 (BFP-ZO-1, cyan). Time interval, 5 s; video frame rate, 20 fps.

**Video 6:** Time-lapse confocal imaging of gastrula-stage *Xenopus laevis* epithelium treated with a mix of 20 µM BAPTA-AM and 100 µM 2-APB for 1 hour. Movie shows severe, long lasting ZO-1 breaks (white arrows) and reduced activation of Rho flares (yellow arrowheads) in the absence of calcium flashes (GCaMP6m). All channels shown in FIRE LUT. Time interval, 5 s; video frame rate, 50 fps. Related to Fig. 3 C.

**Video 7**: Time-lapse confocal imaging shows the increased frequency of Rho flares (mCherry-2xrGBD, grayscale) in embryos treated with 12.5 µM GsMTx4 (green arrowheads) compared to vehicle (yellow arrows). Dividing cells are marked by cyan asterisks. Time interval, 21 s; video frame rate, 20 fps. Related to Fig. 5 C.

**Video 8:** Time-lapse confocal imaging showing leaks (FluoZin-3, orange LUT) in embryos expressing BFP-ZO-1 (FIRE LUT) and active Rho probe (mCherry-2xrGBD, FIRE LUT). Video shows the local increase of FluoZin-3 (white arrow) preceding the Rho flare (yellow arrowhead) at the site of ZO-1 decrease (white arrowhead) in vehicle (water) treated embryos. Note the repeated local increase of FluoZin-3 (green and white arrows) followed by short duration Rho flare (yellow arrowhead) in embryos treated with 12.5 μM GsMTx4. Time interval, 21 s; video frame rate, 5 fps. Related to Fig. 5 D-E.

**Video 9:** Time-lapse confocal imaging shows repeating Rho flare (yellow arrowhead) and F-actin accumulation (yellow arrow) at the site of ZO-1 loss (white arrow) in embryos treated with 12.5 μM GsMTx4 compared to vehicle (water). Note the reduced ZO-1 reinforcement (white arrow) following Rho flare at 410 s and recurring ZO-1 break at the same site at time 1450 s in GsMTx4 treated embryo. Time interval, 10 s; video frame rate, 20 fps.
